# One-day phenotypic drug susceptibility testing for *Mycobacterium tuberculosis* variant *bovis* BCG using single-cell imaging and a deep neural network

**DOI:** 10.1101/2024.05.20.594971

**Authors:** Buu Minh Tran, Jimmy Larsson, Anastasia Grip, Praneeth Karempudi, Johan Elf

## Abstract

Drug-resistant tuberculosis (TB) kills approximately 200,000 people every year. A contributing factor is the slow turnaround time associated with anti-tuberculosis drug susceptibility diagnostics. The prevailing gold standard for phenotypic drug susceptibility testing (pDST) takes at least two weeks. In this study, we used *Mycobacterium tuberculosis* variant *bovis* BCG (*M. bovis* BCG) and *Mycobacterium smegmatis* as models for tuberculous and nontuberculous pathogens. The bacteria were loaded into a microfluidic chip, trapping them in microchambers, and allowing simultaneous tracking of single-cell growth with and without antibiotic exposure. A deep neural network image-segmentation algorithm was employed to quantify the growth rate over time and determine how the strains responded to the drugs compared to the untreated reference. We determined that the response time of the susceptible strains to isoniazid (INH), ethambutol (EMB), and linezolid (LZD) at MIC was within 3 hours and 1.5 hours for *M. bovis* BCG and *M. smegmatis*, respectively. Resistant strains of *M. smegmatis* were identifiable within 3 hours, suggesting that growth-based pDST can be conducted in less than 12 hours for slow-growing *M. bovis* BCG. The results obtained for *M. bovis* BCG are most likely comparable to what we expect for *M. tuberculosis* as these strains share 99.96% genetic identity.

## Introduction

Tuberculosis (TB) has afflicted humans for millennia and remains a major public health threat^1^. As COVID-19 fatalities decrease, *Mycobacterium tuberculosis* (*Mtb*), the causative agent of TB, has reclaimed its position as the primary infectious cause of death worldwide. An estimated 10.6 million people contracted TB in 2022, and in the same year, a total of 1.3 million died from the disease^2^. The approximate number of people who developed TB is the tip of the iceberg; about a quarter of the global population may be infected with TB^3^. A major driver of TB mortality is the increasing prevalence of drug-resistant TB (DR-TB), multidrug-resistant TB (MDR-TB), pre-extensively drug-resistant TB (pre-XDR-TB), and extensively drug-resistant TB (XDR-TB) infections^4–9^. In 2022, nearly half a million people developed multidrug-resistant or rifampicin-resistant TB (MDR/RR-TB)^2,10^. In 2019, around 25% of global deaths from antimicrobial-resistant infections were attributed to rifampicin-resistant TB^10^. Additionally, an estimated 1 million cases of TB resistant to isoniazid (another essential first-line drug, without simultaneous rifampicin resistance) emerged in the same year^10^.

Treating drug-resistant TB requires lengthy, expensive drug regimens and is often linked to worse treatment outcomes, including a higher mortality rate, compared to susceptible cases^11–14^. In a multicentre cohort study, more than half of the misdiagnosed drug-resistant tuberculosis cases received inadequate therapy resulting in death^15^. Therefore, an efficient TB treatment regimen requires a combination of antibiotics to minimize treatment failure due to resistance and the emergence of multidrug-resistant TB. More importantly, it is recommended that drug susceptibility tests (DST) are enforced to guide initial treatment selection^16,17^.

Presently, both genotypic and phenotypic drug susceptibility testing (gDST and pDST) methods are employed in combination to accurately determine the drug susceptibility of *Mtb.* Genotypic methods like nucleic acid amplification tests (NAAT) and line probe assays (LPA), are widely used. The Xpert® MTB/XDR NAAT (Cepheid), which detects resistance to isoniazid (INH), fluoroquinolones (FLQ), ethionamide (ETH), and second-line injectable drugs (SLIDs), can detect 16 resistance mutations with clinical isolate DNA in under 90 minutes^18^. The LPA Genotype MTBDRplus (Hain Lifescience GmBH) can detect RIF- and INH-associated mutations with high specificity in less than 6 hours^19^. Despite the reported short turnaround time of gDST in an optimal research laboratory, the time to result under operational conditions ranges from 1 to 10 working days^9^. Whole genome sequencing (WGS) is also recommended in gDST, with the ability to detect resistance mutations outside the limited spectra of NAAT and LPA^19–21^. Although WGS gDST methods are generally effective for *Mtb*, they are expensive, unfit for point-of-care testing, inaccessible to low-resource areas, problematic for heteroresistant infections, cannot detect novel resistance mutations^22,23^, and can only determine if an isolate is likely to be resistant, not that it is susceptible.

Phenotypic drug susceptibility testing (pDST) using liquid or solid media remains the gold standard DST method. Liquid culture-based pDST such as BACTEC MGIT 960 (Becton Dickinson) produces TB diagnostic results within two weeks, likely the fastest commercially available option in clinical use^9,19^. pDST on solid media is simple and accessible, but the turnaround time is 21-45 days or longer^22^. The long turnaround times for pDST were observed to result in acquired drug resistance, thereby affecting the efficacy of tuberculosis treatment^24^. Despite the limitations arising from the slow growth of *Mtb*, pDST remains dependable and proficient in detecting drug resistance resulting from unidentified mechanisms overlooked by genotypic analysis^9^. A nanomotion technology detecting the vibration of bacterial cells attached to cantilevers has been used for drug susceptibility testing of *Mtb,* providing results in 21 hours^25^. However, this technology requires an expensive instrument locked for one sample and one antibiotic for the whole assay.

A combination of microfluidics and microscopy has recently been demonstrated to speed up growth-dependent pDST^26,27^. With this high-throughput methodology, bacteria are trapped in thousands of microchannels on a microfluidic chip. Different areas of the chip are exposed to different media, making it possible to study the effect of antibiotic drugs compared to untreated control. The setup allows for highly controlled experiments where growth impact is measured from the length extension of individual cells instead of waiting for the cells to multiply. Our previous study focused on uropathogenic *Escherichia coli*. We could produce pDST results of clinical samples in less than 30 minutes, with sensitivity and specificity of 86 - 100% compared to a conventional disc-diffusion^26^. For these tests, low bacterial counts (10^4^ CFU/mL) in a spiked sample were sufficient. In the following study, we tuned the method to detect mixed-species samples (*e.g. E. coli, Klebsiella pneumoniae, Staphylococcus aureus*, and *Pseudomonas aeruginosa*) by combining the pDST with fluorescence *in situ* hybridization to identify each species following susceptibility testing. A deep learning method was also embedded in the analysis pipeline to increase the cell segmentation robustness^27^.

pDST in microfluidic chips for mycobacteria has yet to be widely implemented but some studies on the physiology of mycobacterial cells have been performed using microfluidic devices. Aldridge *et al.*, (2012) loaded *M. smegmatis* into microfluidic chips to study their asymmetric growth at the single-cell level and test the differential susceptibility to antibiotic stress^28^. Wakamoto *et al.*, (2013) also used microfluidic culture and time-lapse microscopy to study the dynamic persistence of *M. smegmatis* under antibiotic stress with isoniazid^29^. Baron *et al.,* (2020) used a microfluidic acoustic-Raman platform to assess the impact of isoniazid on *M. smegmatis* in real time^30^. Another microfluidic device was developed by Wang *et al.,* (2021) to visualize the long-term growth of *M. smegmatis* under antibiotic pressures. *M. tuberculosis* was loaded into a microfluidic chip by Chung *et al.,* (2023) to investigate the growth at the single-cell level^31^. Notably, microfluidics and direct imaging combined with agarose channels were used for rapid pDST on various bacterial strains, including *Mtb*^32–35^. The system, called QuantaMatrix microfluidic agarose channel (QMAC), was integrated with MGIT liquid culture, resulting in a short turnaround time of the culture identification (∼2 weeks) and subsequent pDST from *Mtb* clumps (∼1 week)^35^.

In this study, we present a rapid pDST approach for *M. bovis* BCG and *M. smegmatis* as models for slow-growing tuberculous and fast-growing nontuberculous pathogens. The approach combines microfluidics, single-cell imaging, and deep neural network (DNN)-based image analysis.

## Results

### Fast phenotypic drug susceptibility testing (pDST) on a microfluidic chip

The workflow for the fast pDST for *M. bovis* BCG is shown in Fig. 1a and the details are given in the Supporting Information (SI). We start with a liquid culture sample. The cells in the liquid culture are loaded into the microfluidic device for compartmentalization in microchambers. The growth of the cells in the microchambers under different antibiotic exposures is monitored by phase-contrast microscopy. The resulting image data is segmented with a DNN-based method.

**Figure 1.**
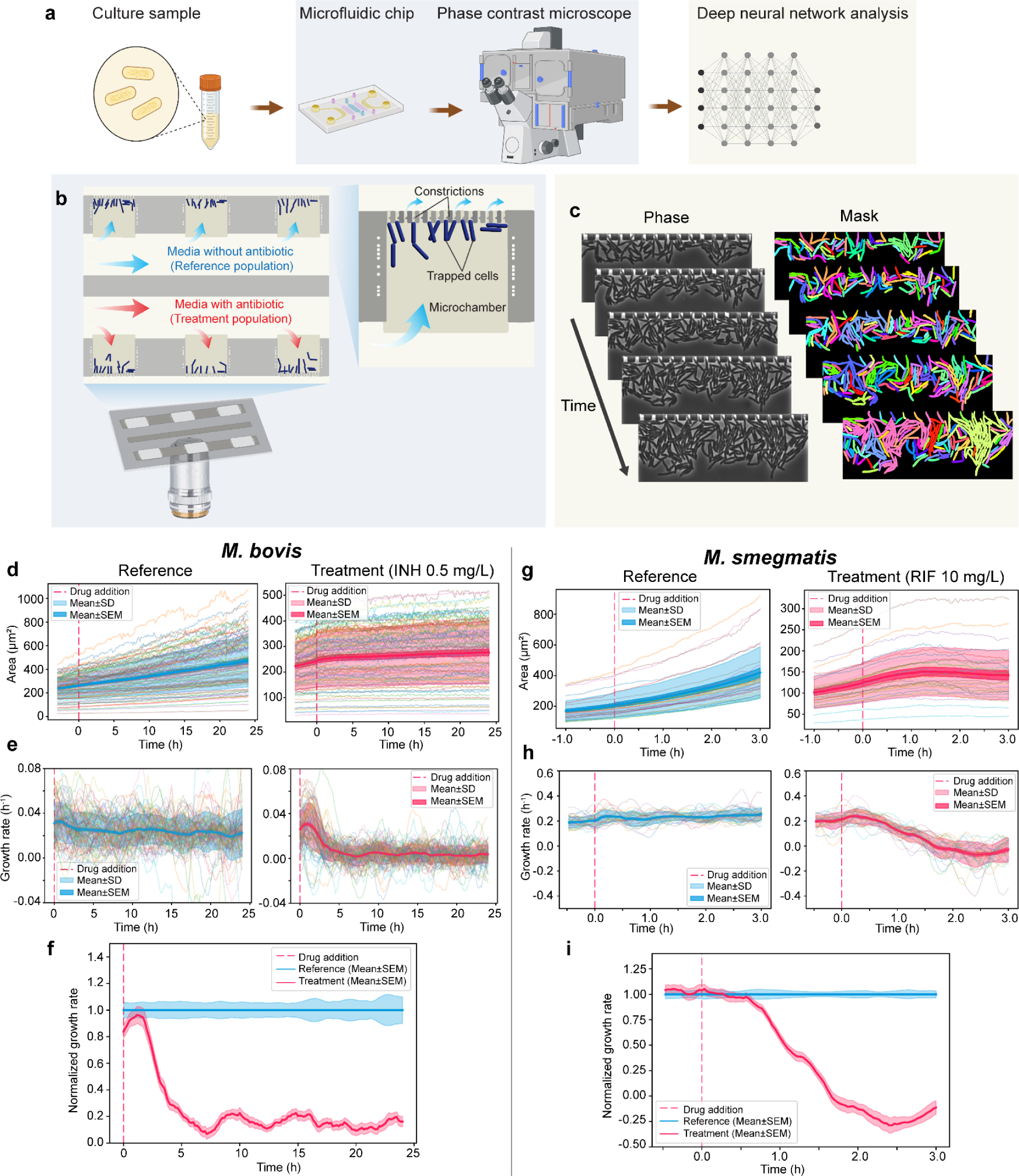
Schematic representation of the workflow of fast pDST for *M. bovis* BCG and *M. smegmatis.* **a**. Main steps in the workflow starting from a culture sample, compartmentalization of cells in a microfluidic chip, time-lapse phase contrast microscope, and DNN image analysis. **b**. Illustration of the microchamber cell traps. **c**. Analysis of time-lapse stacks using the DNN called Omnipose for cell segmentation. **d – i.** Step-by-step growth rate calculation and normalization for *M. bovis* BCG (**d**, **e**, and **f**) and *M. smegmatis* (**g**, **h**, and **i**). **d**. Total areas of *M. bovis* BCG cells are plotted as a function of time from the reference population (left) and treatment population (right; INH 0.5 mg/L). The interval between frames is 10 min, and each line is data from an individual microchamber. The dashed red line indicates the time when the drug is added to the media of the treatment. **e**. Growth rates are estimated with a sliding window time of 3 hours for *M. bovis* BCG. **f**. The growth rate normalization of the treatment and reference population is the pDST profile of *M. bovis* BCG in INH 0.5 mg/L (mean±SEM). **g**, **h**, and **i**. Corresponding area growth (**g**) growth rate (**h**) of the reference (left) and treatment population (right: RIF 10 mg/L) of *M. smegmatis;* and the pDST profile of *M. smegmatis* in RIF 10 mg/L is shown by normalized growth rates in (**i**). Measurements of *M. smegmatis* were carried out at 2 min intervals and growth rates were calculated with a sliding window time of 30 min. Fig. 1a is created using www.biorender.com.

Fig. 1b shows the microfluidic chip design. The chip has two rows of microchambers that function as cell traps; there are 100 microchambers with 50 × 60 × 1 µm dimensions on each row. At the end of each microchamber, constrictions (300 nm) prevent the cells from escaping into the back channel while allowing a constant media flow over the cells. Once the cells have been loaded into the microchambers, their growth is monitored by phase-contrast time-lapse imaging. To find the outlines of single cells in each image frame, we use the morphology-independent segmentation algorithm of Omnipose^36^ (Fig. 1c).

We describe the growth rate estimation and normalization details in Fig. 1d to Fig. 1i. In this work, *M. bovis* BCG and *M. smegmatis* are used as models for tuberculous and nontuberculous pathogens. Due to the difference in their growth rate, we applied the measurement time of 3 + 24 hours and 1 + 3 hours for *M. bovis* BCG (Fig. 1d, 1e, and 1f) and *M. smegmatis* (Fig. 1g, 1h, and 1i) respectively; the former terms are the times when media without drugs are supplied to both rows of microchambers, and the latter terms are the times when the drug is supplied to the medium in the treatment row. The expansion of the total cell area was monitored for 27 hours (*M. bovis BCG)* and 4 hours (*M. smegmatis*), indicating the cells’ growth within the microchambers. Fig. 1d shows the total areas of *M. bovis* BCG plotted as a function of time for the reference population (left) and treatment population (right; INH 0.5 mg/L). Each line in Fig. 1d presents data from an individual microchamber. Images were captured every 10 min. The drug was added to the treatment population after 3 hours (dashed red line). The growth rates were estimated with a sliding window of 3 hours for *M. bovis* BCG, and thus, the growth rates are plotted from 3 hours after starting the experiment (Fig. 1e). Fig. 1f shows the growth rate of *M. bovis* BCG in INH 0.5 mg/L normalized to the reference population. This type of normalization has previously been used^26,27^ to reduce the isolate dependence in the antibiotic response. The area growth and growth rates of the reference (left) and treatment population (right: RIF 10 mg/L) of *M. smegmatis* are shown in Fig. 1g and 1h. The pDST profile of *M. smegmatis* in RIF 10 mg/L is shown by normalized growth rates in Fig. 1i. In the case of *M. smegmatis*, we measured at 2 min intervals and growth rates were calculated with a sliding window time of 30 min. The occasional negative growth rate indicates shrinkage of the cell area and can for example be due to lysis.

### Data analysis using a deep neural network

Accurate cell segmentation tools for mycobacteria are not available. Optical single-cell studies on mycobacteria have so far been performed using manual annotation, likely due to their small and varied cell size and the abnormally asymmetric aging and division^28,31^. An artificial neural network architecture, U-net^37^, was used for the automatic segmentation of TB cords from lens-free microscopy images^38^. Similar to the U-net, the general segmentation algorithm, Omnipose^36^, generates cell outlines using intermediate fields learned by a convolutional neural network, thereby facilitating bacterial cell segmentation. Omnipose achieved high accuracy and morphological independence in the segmentation of mixed bacterial samples, antibiotic-treated, and elongated or branched cells^27,36^.

We created a 120-image ground truth training dataset of mycobacterial cells under diverse experimental conditions, encompassing *M. bovis* BCG and *M. smegmatis* captured by two different cameras (Methods) on agarose pads and in microchambers, and cells subjected to antibiotic treatment. High-density mycobacterial clumps or cords in microchambers were membrane stained using 3HC-3-Tre dye^39^ to assist the annotation (Fig. 2a). The models Mycobact_1 to Mycobact_4 were trained from scratch while Mycobact_5 and Mycobact_6 were fine-tuned based on the Omnipose default model for bacteria in phase contrast. The training parameters are shown in SI Table 1. The segmentation performance of the new models was evaluated by matching the segmented masks to the ground truth masks at various matching precision thresholds. Quantitatively, we assessed the model performance by comparing their average Jaccard Index^40^ as a function of the intersection over union (IoU) threshold on the same dataset (Fig. 2b and 2c). IoU scores range from zero to one, with scores greater than 0.8 indicating when masks are visually indistinguishable from ground truth according to human experts^36^. In Fig. 2b and 2c, the performance of all new models did not remarkably vary with the IoU threshold less than 0.75; with the IoU threshold greater than 0.75, the training-from-scratch models outperformed the fine-tuned models. The difference in performance suggests that the images of mycobacteria are unique datasets or the pre-trained Omnipose model might carry inherent biases against mycobacterial data. It is noted that the general performance evaluation used an image dataset from two cameras with different pixel sizes, owing to our experiments conducted on two different microscopes. However, the performance of the newly trained model significantly improved when using images from a single camera (Fig. 2b, Mycobact_2*). This improvement can be attributed to the use of more accurate parameters for the images from one camera, such as diameter and minimal cell size.

**Figure 2.**
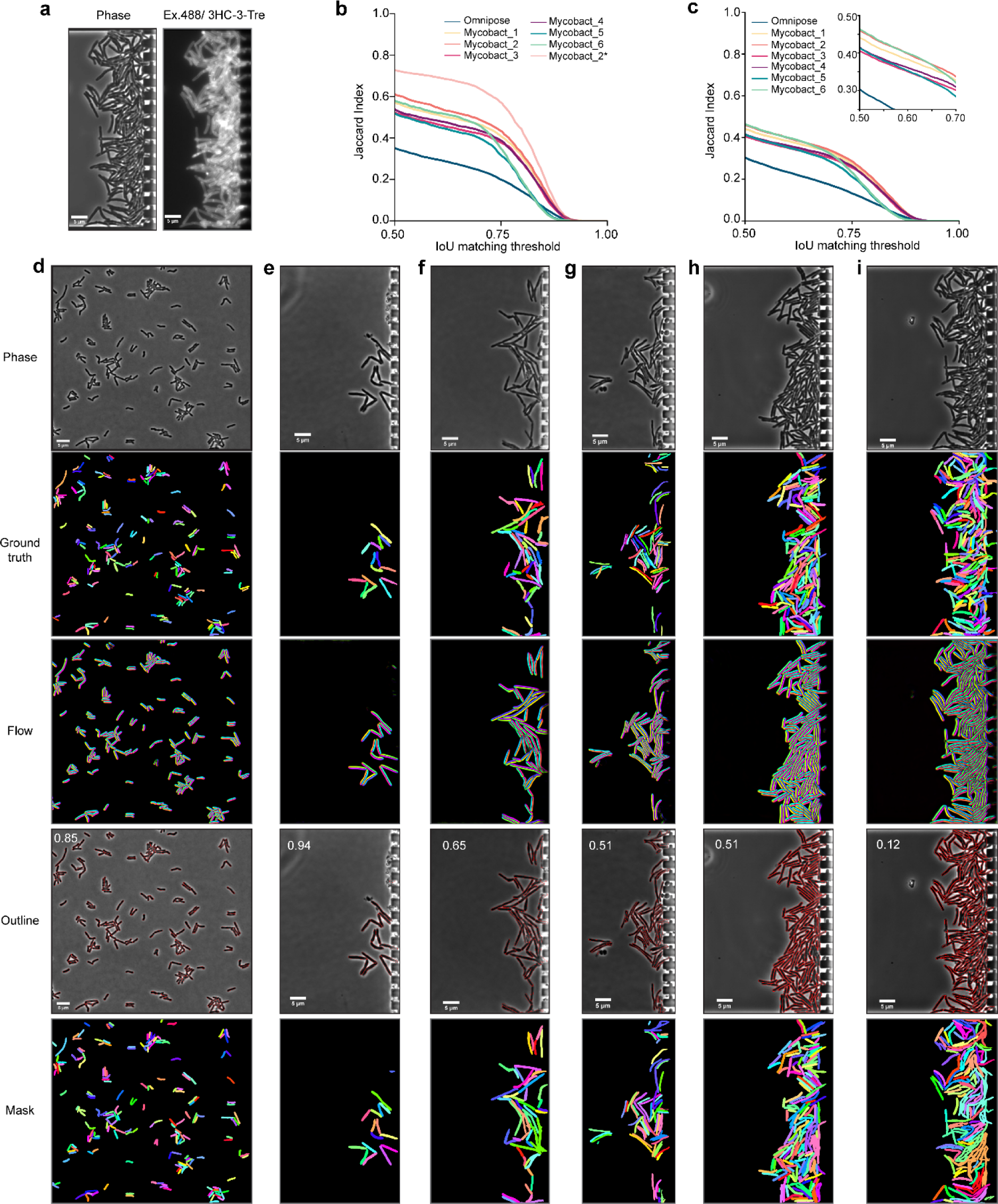
Training new models for mycobacterial cell segmentation. **a**. Phase contrast image and the corresponding image of cell membrane stained by 3HC-3-Tre dye for labeling assistance in high-density cell microchambers. **b-c**. Segmentation performance of different training and fine-tuning parameters using data (**b**) excluding and (**c**) including high-density cell microchambers. Mycobact_2* in **(b)** is the Mycobact_2 model with the segmentation performed using images from one camera. Inset in **(c)** is a zoom-in at a smaller scale in the x and y-axis. **d-i**. Representative micrographs of mycobact_2 model for different mycobacterial cells in various conditions – (**d**) Flat and well-separated *M. smegmatis* on agarose gel, (**e**) and (**f**) Separated *M. smegmatis* in the microfluidic chamber, (**g**) *M. bovis* BCG in the microchamber, (**h**) and (**i**) Relatively and highly dense *M. smegmatis* cells. Ground truth mask is manually labeled from phase images. The neural network produces the flow field as an intermediate output for outline mask reconstruction. Average precision at an IoU threshold of 0.5 (AP@0.5) for the entire image is reported on the top left corner of the outline—scale bar 5 µm.

**Table 1.** Training/fine-tuning parameters of models for mycobacteria using Omnipose network ^36^.

| No. | Model | Training parameters |
| --- | --- | --- |
| 1. | Omnipose | default Omnipose model for bacteria phase contrast (bact_omnitorch_0) |
| 2. | Mycobact_1 | trained from scratch, 4000 epochs, nclasses=2, learning_rate=0.1, 240 images |
| 3. | Mycobact_2 | trained from scratch, 4000 epochs, nclasses=3, learning_rate=0.05, 240 images |
| 4. | Mycobact_3 | trained from scratch, 4000 epochs, nclasses=3, learning_rate=0.05, 738 images |
| 5. | Mycobact_4 | trained from scratch, 8000 epochs, nclasses=3, learning_rate=0.01, 738 images |
| 6. | Mycobact_5 | fine-tuned from default Omnipose model for bacteria phase contrast, 1300 epochs, nclasses=3, learning_rate=0.1, 240 images |
| 7. | Mycobact_6 | fine-tuned default Omnipose model for bacteria phase contrast, 4000 epochs, nclasses=3, learning_rate=0.1, 240 images |

Overall, the segmentation performance of the new models showed remarkable improvements compared to the default bacteria model from Omnipose, particularly on low cell density data (Fig. 2b and Fig. 2d - 2g). High cell density data (*e.g*. clumps or cords) were problematic due to overlapping cells and difficulty distinguishing single-cell boundaries. Instead of identifying single cells, the entire blob of cells was segmented as one (Fig 2c and Fig.2h - 2i). Movies showing image segmentation performance are supplied in the <u>SI</u>.

### Fast detection of response to drug treatment

We used the fast pDST to determine the response of *M. bovis* BCG and *M. smegmatis* (NCTC 8159 and mc^2^ 155) to three first-line drugs, rifampicin (RIF), isoniazid (INH), and ethambutol (EMB), and the third-line drug linezolid (LZD). MIC values of *M. smegmatis* and *M. bovis* BCG were determined using the microplate-based Alamar Blue assay (MABA)^41,42^ and EUCAST broth microdilution assay (BMDA)^43^, respectively. Fig. 3a - 3d show the pDST profiles of *M. bovis* BCG tested at the MICs of RIF (0.06 mg/L), INH (0.5 mg/L), EMB (2.5 mg/L), and LZD (0.5 mg/L). We could measure a difference in growth rates between the treated and control populations under 3 hours for INH, EMB, and LZD and under 12 hours for RIF. Given that *M. bovis* BCG has a replication time of 18 - 24 hours^44^, a difference in growth rate between the treatment and reference population could be detected at ⅛ to ⅙ of a generation time for slow-growing tuberculous mycobacteria (*i.e. M. bovis* BCG). For fast-growing mycobacteria, the difference in growth rates between the two populations could be detected in under 1 hour in all cases except for *M. smegmatis* mc^2^ 155 in LZD 0.16 mg/L where the difference was detected after about 1.5 hours (Fig. 3e - 3l). The average doubling time of *M. smegmatis* is 3 hours^28,45,46^, which means that the growth-rate difference could be detected after ⅓ of a generation. In Supplementary Figure 1, we show repeats of these experiments. The response curves are reproducible for the same antibiotic, but the shapes of the curves differ for different antibiotics, implying that the response time is limited by the biology rather than the measurement.

**Figure 3.**
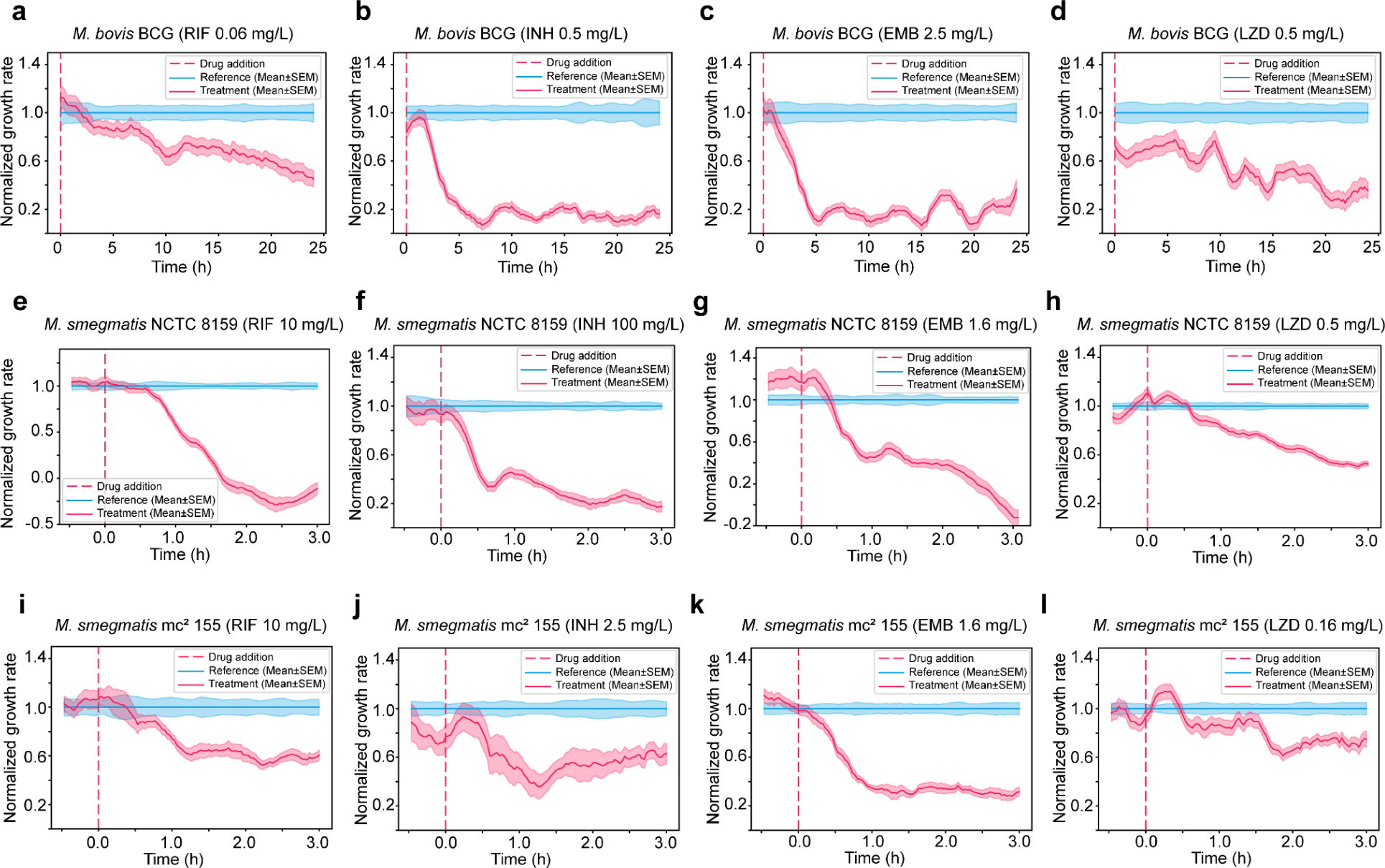
Fast detection of response to drug treatment. pDST assays detecting the fast response of *M. bovis* BCG to (**a**) rifampicin (RIF), (**b**) isoniazid (INH), (**c**) ethambutol (EMB), and (**d**) linezolid (LZD); of *M. smegmatis* NCTC 8159 to (**e**) RIF, (**f**) INH, (**g**) EMB, and (**h**) LZD; and *M. smegmatis* mc^2^ 155 to (**i**) RIF, (**j**) INH, (**k**) EMB, and (**l**) LZD.

### Detection of drug-resistant strains

We carried out the fast pDST assay on drug-resistant strains to get a step closer to clinically relevant conditions. This is crucial because resistant strains vary in pDST profiles for different drugs. In the presence of antibiotics, the resistant strains could respond similarly to the susceptible strains at the beginning of the measurement but recover the growth rate later^29,47–49^. We thus have to make sure that we image the cells for a sufficiently long time to capture this growth recovery phase. For this reason, we conducted the pDST on RIF, INH, and LZD-resistant strains (Fig. 4), which were laboratory-evolved and characterized^50^. These strains harbored genomic mutations in the genes of *rpoB* (RIF resistance), *inhA* and *katG* (INH resistance), and *oxiR* (LZD resistance)^50^. The pDST profiles of the drug-resistant strains were compared with the susceptible parental strain *M. smegmatis* NCTC 8159 (Fig. 3e, 3f, and 3h).

**Figure 4.**
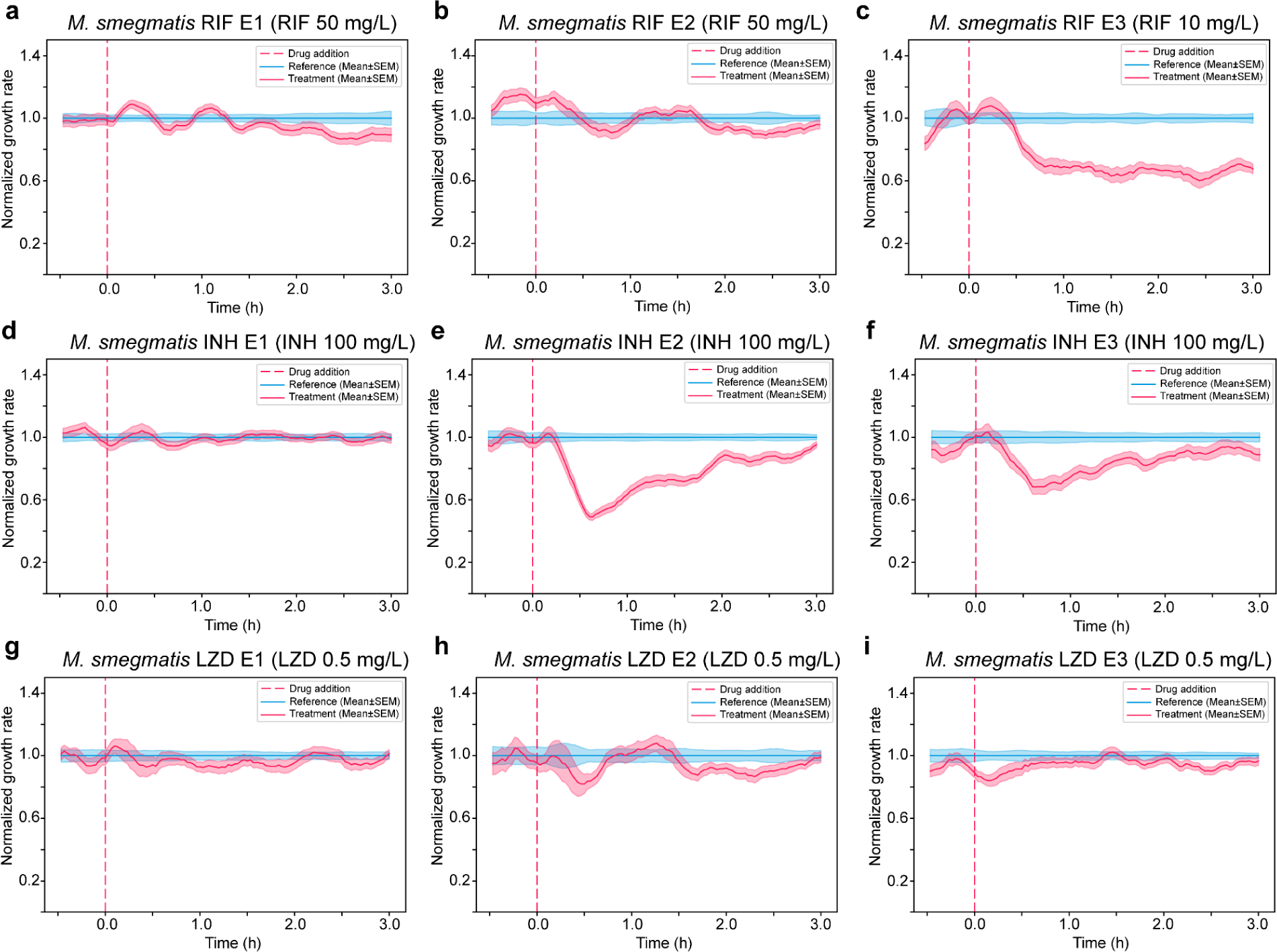
Fast detection of resistant strains. pDST profiles of lab-evolved resistant strains derived from *M. smegmatis* NCTC 8159 including (**a**) *M. smegmatis* RIF E1, (**b**) *M. smegmatis* RIF E2, (**c**) *M. smegmatis* RIF E3 in rifampicin treatment, (**d**) *M. smegmatis* INH E1, (**e**) *M. smegmatis* INH E2, (**f**) *M. smegmatis* INH E3 in isoniazid treatment, and (**g**) *M. smegmatis* LZD E1, (**h**) *M. smegmatis* LZD E2, (**i**) *M. smegmatis* LZD E3 in linezolid treatment.

Fig. 4a to 4c show the pDST profiles of *M. smegmatis* RIF E1 to E3. The growth of the treatment population of RIF E1 and E2 strains is comparable with the reference population at 50 mg/L RIF concentration (Fig. 4a and 4b). RIF E3 is more sensitive than RIF E1 and E2 even at lower RIF concentration (10 mg/L) (Fig. 4c), but more resistant compared to the wild-type strain at the same concentration of RIF (Fig. 3e). These results from the 3-hour pDST assay indicate that RIF E3 is less resistant than RIF E1 and E2, in agreement with the 48-hour REMA assay (SI Table 2). For INH-resistant strains, the growth of the treatment population of *M. smegmatis* INH E1 is indistinguishable from the reference population implying high resistance (Fig. 4d). INH E2 and INH E3 responded to INH treatment similarly; the growth rates dropped dramatically in the first 30 min after the drug was added and then gradually recovered reaching the reference level in 3 hours (Fig. 4e and 4f). At 0.5 mg/L LZD concentration, the growth rates of the three LZD-resistant strains were not greatly affected by the antibiotic (Fig 4g - 4i) as opposed to the susceptible parental strain (Fig. 3h). We show repeats of these experiments with resistant strains in Supplementary Figure 2. Overall, each resistant strain reacted differently to the corresponding drug treatment, but the resistance could in all cases be detected within 3 hours by the fast pDST assay.

**Table 2.** MIC values of the tested strains were determined using the resazurin microtiter assay (REMA)^41^ and EUCAST broth microdilution assay^43^

| No. | Strain | MIC |  |  |  | Remarks |
| --- | --- | --- | --- | --- | --- | --- |
|  |  | RIF | INH | EMB | LZD |  |
| 1. | *M. bovis* BCG WT | 0.06 | 0.5 | 2 | 0.5 | ATCC - 35734 |
| 2. | *M. smegmatis* mc<sup>2</sup> 155 WT | 10 | 2.5 | 1.6 | 0.16 | Leif Kirsebom |
| 3. | *M. smegmatis* NCTC 8159 WT | 10 | 50 | 1.6 | 0.4 | 50 |
| 4. | *M. smegmatis* NCTC 8159 RIF E1 | >100 (50*) |  |  |  |  |
| 5. | *M. smegmatis* NCTC 8159 RIF E2 | >100 (50) |  |  |  |  |
| 6. | *M. smegmatis* NCTC 8159 RIF E3 | 50 (50) |  |  |  |  |
| 7. | *M. smegmatis* NCTC 8159 INH E1 |  | >100 (100) |  |  |  |
| 8. | *M. smegmatis* NCTC 8159 INH E2 |  | >100 (100) |  |  |  |
| 9. | *M. smegmatis* NCTC 8159 INH E3 |  | >100 (100) |  |  |  |
| 10. | *M. smegmatis* NCTC 8159 LZD E1 |  |  |  | 50 (0.5) |  |
| 11. | *M. smegmatis* NCTC 8159 LZD E2 |  |  |  | 12.5 (0.5) |  |
| 12. | *M. smegmatis* NCTC 8159 LZD E3 |  |  |  | 50 (0.5) |  |

(\*) Values in the parentheses are the lab-evolved selective concentrations<sup>50</sup>

## Discussion

The World Health Organization recommends drug susceptibility testing (DST) for samples from all individuals exhibiting clinical symptoms of active TB, despite the existing tools being inadequate to achieve this objective^22,51–53^. Rapid and accurate results from DST of clinical TB samples are vital to avoid misprescribing ineffective or toxic additional drugs and control the spread of resistant variants. Furthermore, the optimal treatment of each tuberculosis patient can be evaluated and implemented more rapidly, shortening treatment regimens^19^. Culture-based phenotypic DST (pDST) methods for anti-TB drugs are reliable and reproducible, yet time-consuming, demanding sophisticated lab setups, trained personnel, and rigorous quality control^53^. The pDST presented in this study for slow-growing mycobacteria is inspired by the fast antibiotic susceptibility test (fASTest) developed in our laboratory for UTI pathogens^26^. In this study, we modified the chip to accommodate mycobacterial cells and robustly analyzed the image data using a deep neural network.

In the current workflow, we would start a diagnostic assay after a culture-positive TB diagnosis. This avoids the complications of sample prep from sputum and means that we could load relatively high concentrations of exponentially growing bacteria. We currently use 5×10^4^ to 1×10^5^ CFU/mL, which could be pushed down if needed. Another aspect of working with a culture-positive sample is that we would only replace the current two-week pDST, thus not radically changing the TB diagnosis workflow. By applying fast pDST, significant deviations in growth rates between the treatment and reference (control) bacterial populations at MICs of INH, EMB, and LZD are observable within 3 hours for slow-growing tuberculous mycobacteria, with a generation time of 18 to 24 hours. This contrast in growth is discernible after approximately 1 - 1.5 hours for the fast-growing *M. smegmatis.* We have not yet been able to test the assay on resistant strains of *M. bovis* BCG. However, based on our observations that resistant *M. smegmatis* can be identified in twice the time it takes to observe the antibiotic impact in the susceptible strain, we estimate that resistance to INH, EMB, and LZD can be determined in as little as 6 hours in *M. bovis* BCG. The time may be shortened further by running the assay at concentrations higher than the MIC.

Our fast pDST can help reduce turnaround time (TAT), allowing for more effective drug response monitoring, which benefits patients. Doctors could choose different drug combinations or adjust the dose accordingly. Moreover, the short TAT of the assay minimizes the acquired resistance developed during the susceptibility testing^24^.

Fast and sensitive methods for TB drug susceptibility testing using direct microscopic observation of broth cultures have been proposed before. However, the median turnaround time for these tests was 5.4 to 9.5 days^35,54–56^. In our method an accurate growth rate is calculated for thousands of single cells, allowing us to reach statistical significance for growth-rate differences quicker than what would be possible for the corresponding bulk measurement. The fact that different antibiotics display different response curves implies that we are looking at biological differences and it is unlikely that any phenotypic method could be faster.

This work presents the proof-of-principle of a rapid pDST for mycobacteria, particularly the slow-growing species. In the current setup, we implemented the test utilizing a state-of-the-art research microscope followed by subsequent processing of the acquired imaging data. We also used *M. bovis* BCG as a slow-growing tuberculous mycobacterial model because it requires a lower level of biosafety. To take the system one step closer to the clinic, fluid automation in a closed system should be implemented and tested for highly virulent and resistant *M. tuberculosis* from clinical samples.

## Materials and methods

### Microfluidic chip design and fabrication

The microfluidic chip is composed of a micro-molded silicon elastomer layer (polydimethylsiloxane (PDMS); Sylgard 184) covalently bonded with a borosilicate cover glass (thickness no. 1.5, 22×40 mm, VWR). Details of the microfluidic chip design were described in the previous work^26,57^, and microchannels were replaced by microchambers (Fig. 1b). The numbering of the ports for the microfluidic chip was illustrated in these references. Tubing (VWR, TYGON VERNAAD04103) was connected to the chip via a metal tubing connector. We used ports 5.1, 5.2, and 6.0 to maintain back-channel pressure, port 2.0 to load cells, and ports 2.1 and 2.2 to supply growth media with and without drugs. Flow control was regulated by the OB1-Mk3 regulator (Elveflow).

### Mycobacterial strains and antibiotics

*Mycobacterium tuberculosis* variants *bovis* BCG (ATCC 35734) and *Mycobacterium smegmatis* (mc^2^ 155 and NCTC 8159) were used as models for tuberculous and nontuberculous pathogens. *M. smegmatis* mc^2^ 155 WT was kindly provided by Leif Kirsebom. *M. smegmatis* NCTC 8159 WT and laboratory-evolved resistant strains (RIF E1, RIF E2, RIF E3, INH E1, INH E2, INH E3, LZD E1, LZD E2, LZD E3) was kindly given by Tomoya Maeda^50^. Antibiotics (rifampicin (RIF, R3501), isoniazid (INH, I3377), ethambutol (EMB, E4630) and linezolid (LZD, PHR1885) were purchased from Sigma-Aldrich. Stock solutions were prepared by dissolving active agents in DMSO at 10,240 mg/L and stored at -20°C.

### Media and cultural conditions

We used Middlebrook 7H9 (Sigma-Alrich M0178) liquid medium as growth medium (GM) supplied with 10% Acid-Dextrose-Catalase (ADC) solution, 0.005% glycerol (w/v; Sigma-Aldrich G5516), 0.0005% Tween 80 (w/v; Sigma-Aldrich S6760), and 0.17% Pluronic F-108 (w/v; Sigma-Aldrich 542342). ADC solution contains 8.5 g NaCl (Sigma-Aldrich S3014), 50 g bovine serum albumin (Sigma-Aldrich A2153), 20 g dextrose (Sigma-Aldrich D9434), 0.03 g catalase (Sigma-Aldrich C9322) per liter. *M. bovis* BCG from glycerol stocks was streaked on egg-based Löwenstein-Jensen (LJ, Sigma-Aldrich 63237-500G-F) solid plates. *M. smegmatis* was grown on LB agar plates. For off-chip culture, overnight growth was prepared by inoculating three colonies into 4 mL GM in round-bottom clear polystyrene culture tubes (VWR, 734-0435) and incubated at 37°C with 200 rpm shaking for 18 hours and 6 days for *M. smegmatis* and *M. bovis* BCG respectively.

### MIC determination for M. bovis BCG

The MIC measurements for slow-growing *M. bovis* BCG were determined using broth microdilution assay (BMDA) following the guidelines from EUCAST^43^. The guideline was for *M. tuberculosis* but adapted for *M. bovis* BCG. First is the preparation of broth and anti-tuberculous drugs. We used 96-well flat-bottom-shaped polystyrene plates (Sigma-Aldrich, Costar 3370). The plates with broth and drugs were prepared and used immediately. The broth was the growth media (GM) for both off-chip and on-chip experiments. A 4X working solution was two-step diluted in GM from an aliquot of a stock solution. Except for the peripheral wells which would be filled with sterile distilled water to prevent desiccation during incubation, 0.1 mL of GM was added to all wells. Subsequently, 0.1 mL of the 4X working solution was added to the wells corresponding to the highest concentration of each drug; Ensure no drug was added to the negative and growth control (GC) wells. A multichannel pipette was used to make 1:2 dilutions by adding 0.1 mL of the antibiotic solution present in the highest concentration row to the following row and discarding the last 0.1 mL of the last row/wells. The second step is the inoculation of culture from the stationary phase of *M. bovis* BCG. The turbidity of the culture was adjusted to McFarland 0.5. Further dilutions of two concentrations at 1:100 and 1:10000 in GM broth, which corresponded to 100% and 1% growth control (GC100% and GC1%), were conducted by tenfold dilution steps. A volume of 0.1 mL of GC100% was added to the wells containing drugs that were prepared in the first step. GC100% and GC1% were also added to designated wells on the plate. The third step is the incubation and MIC determination. After inoculation, a breathable membrane (Sigma-Aldrich, Breathe-Easy® Z380059) was used to seal the plate to avoid drying of the cultures but still kept the air going through, and incubated at 37°C±1°C under normal shaking conditions in Tecan (Sunrise^TM^, Tecan Nordic AB) incubator. Optical density values were read every 1 hour. The negative control should show no growth for the test to be valid. The GC1% should show visible growth and be slower than GC100%. MIC was determined as the lowest concentration of the drug where no visible growth was observed. If there was still insufficient growth of the GC1% after 14 days, incubate until a maximum of 21 days.

### MIC determination for M. smegmatis

MIC values for *M. smegmatis* were determined using the microplate-based Alamar Blue assay (MABA)^41,42^. In short, wells on 96-well microtitre plates were filled with 50 μL of growth medium (GM). Double the required antibiotic concentration was prepared and 100 μl volumes were added to the first column. This was serially diluted to half the concentration by mixing with media only in the subsequent wells till the last second column. The last column was a control without antibiotics. The *M. smegmatis* strains were grown in replicates in GM to an optical density (OD 600nm) of 0.6, diluted to McFarland 0.5, and then further diluted 100-fold. 50 μL of the diluted culture was added to each well such that the final concentration of antibiotic in the first well came down to the desired concentration. The plate was sealed with a breathable membrane (Sigma-Aldrich, Breathe-Easy® Z380059) and then incubated under normal shaking conditions in Tecan (Sunrise^TM^, Tecan Nordic) incubator at 37°C for 40 hours. After 40 hours of incubation, 30 μL of resazurin dye ((Sigma-Aldrich R7017; filter sterilized, 0.2 mg/mL concentration) was added to each well and incubated for 6 hours under shaking and then imaged. MICs were determined as the values of the first well showing no growth as indicated by resazurin dye staining.

### Microfluidic experiments

Mycobacteria cells were loaded after diluting 1:100 of a McFarland 0.5 culture, approximately from 5×10^4^ to 5×10^5^ CFU/mL^43^. We used a Nikon Ti-E and Nikon Ti2-E inverted microscope equipped with CFI Plan Apochromat DM Lambda 100X oil immersion objectives (Nikon) for imaging *M. bovis* BCG and *M. smegmatis*, respectively. Images of *M. bovis* BCG were captured by the Imaging Source (DMK38UX304) camera and *M. smegmatis* by Sona 4.2B-11 (Andor). The microscopes were controlled by Micro-Manager^58^ and in-house built plugins with the optical setup described previously^26,27,57^. The temperature of the microscope stage and the microfluidic chip was maintained at 37°C using a climate enclosure (Oklab). Because of the growth rate differences between *M. bovis* BCG and *M. smegmatis,* the measurement timeline of 3 + 24 hours and 1 + 3 hours were applied for *M. bovis* BCG and *M. smegmatis,* respectively. The former parts were the duration without the addition of drugs on both rows (reference and treatment), and the latter part was the duration of drugs added to the media on the row of treatment. The spatial growth indicated by the total cell area in a microchamber was captured throughout the measurement with the interval time of 10 min and 2 min for *M. bovis* BCG and *M. smegmatis,* respectively. About 30 - 50 microchambers of reference and treatment were monitored simultaneously. The instantaneous growth rates were determined with a sliding window time of 3 hours for *M. bovis* BCG and 30 min for *M. smegmatis*.

### Image processing

The image data were pre-processed for correcting stage shifts and cropping microchambers as regions of interest using an in-house algorithm developed in MATLAB^26^. Empty microchambers were discarded in further analysis. Cropped phase-contrast images of mycobacterial cells in microchambers were segmented using an image-segmentation algorithm called Omnipose^36^. We trained new models for image data of mycobacteria. The training dataset was created with 120 images of manually labelled ground-truth of mycobacterial cells in various experimental conditions including *M. bovis* BCG and *M. smegmatis* on agarose pads and in microchambers, and subjected to antibiotic treatment. Manual labeling ground truth for the training dataset was based on the previous work^27^ using LabelsToROI tools and custom scripts. The membrane of cells in high-density microchambers was stained using 3HC-3-Tre to assist in the labeling^39^. Four models were trained from scratch and two models were fine-tuned from the default Omnipose model for bacterial phase contrast images. We show the training parameters in the SI Table 1. The segmentation performance of new models was quantitatively evaluated by comparing their average Jaccard Index as a function of intersection over union (IoU) threshold on the same dataset.

### Data analysis

The calculation of the growth rate of cells in individual microchambers was completed, as described previously^26,27^, by applying a sliding window of data points (area) and fitting an exponential function: *y* = *ae^bt^*, where y is the total area of segmented cells in a microchamber, *a* is the constant, and b is the growth rate. Sliding windows of *M. bovis* BCG and *M. smegmatis* were 3 hours and 30 min, respectively. The standard error of the mean (SEM) takes the growth rate standard deviation (SD) and divides it by the square root of the total microchamber number. The growth rate normalization was calculated by dividing the mean growth rate of the treatment population by the mean growth rate of the reference population at each time interval. The SEM values for both the reference and treatment were normalized by dividing each SEM value at different time intervals by the mean growth rate of the reference population. The separation time of the treatment population from the reference population was based on the time at the separation of the SEM values between the two populations in the growth rate normalization. For long-term experiments with *M. bovis* BCG, positions losing focus would create outliers in the area growth curve. We removed the outliers before calculating the growth rate, by replacing measurements that deviated more than 5% from the curve fitted in the sliding window by the fitted value. After one round of outlier replacement based on the initial curve fitting, the remaining points were refitted to obtain the reported growth rate.

### Statistics and reproducibility

The study is not randomized. The reference and treatment were measured on the same microfluidic chip in the same set-up and segmentation model for each measurement. Empty microchambers and microchambers with segmentation errors were not included in the calculation of the growth rate. All pDST profiles display normalized growth rate ± SEM. At least two experimental replicates were conducted for each condition.

## Data and code availability

Raw microscopy data, models for segmentation, and analysis code will be publicly available upon final publication.

## Acknowledgement

We acknowledge funding from the Swedish Foundation for Strategic Research (SSF), Knut and Alice Wallenberg Foundation, and Novo Nordisk. We thank Irmeli Barkefors for her helpful comments on the manuscript, and Tomoya Maeda and Leif Kirsebom for providing *M. smegmatis* strains.

## Author contributions

J.E. conceived the pDST method for mycobacteria and supervised the project. B.M.T. performed pDST of *M. bovis* BCG and all data analysis. J.L. performed pDST of *M. smegmatis* NCTC 8159 strains. A.G. performed pDST of *M. smegmatis* mc^2^ 155. P.K. trained mycobact_1 to mycobact_4 models and wrote scripts for model evaluation. J.E. and B.M.T. wrote the paper with input from all authors.

## Supporting Information

**Supplementary Figure 1.**
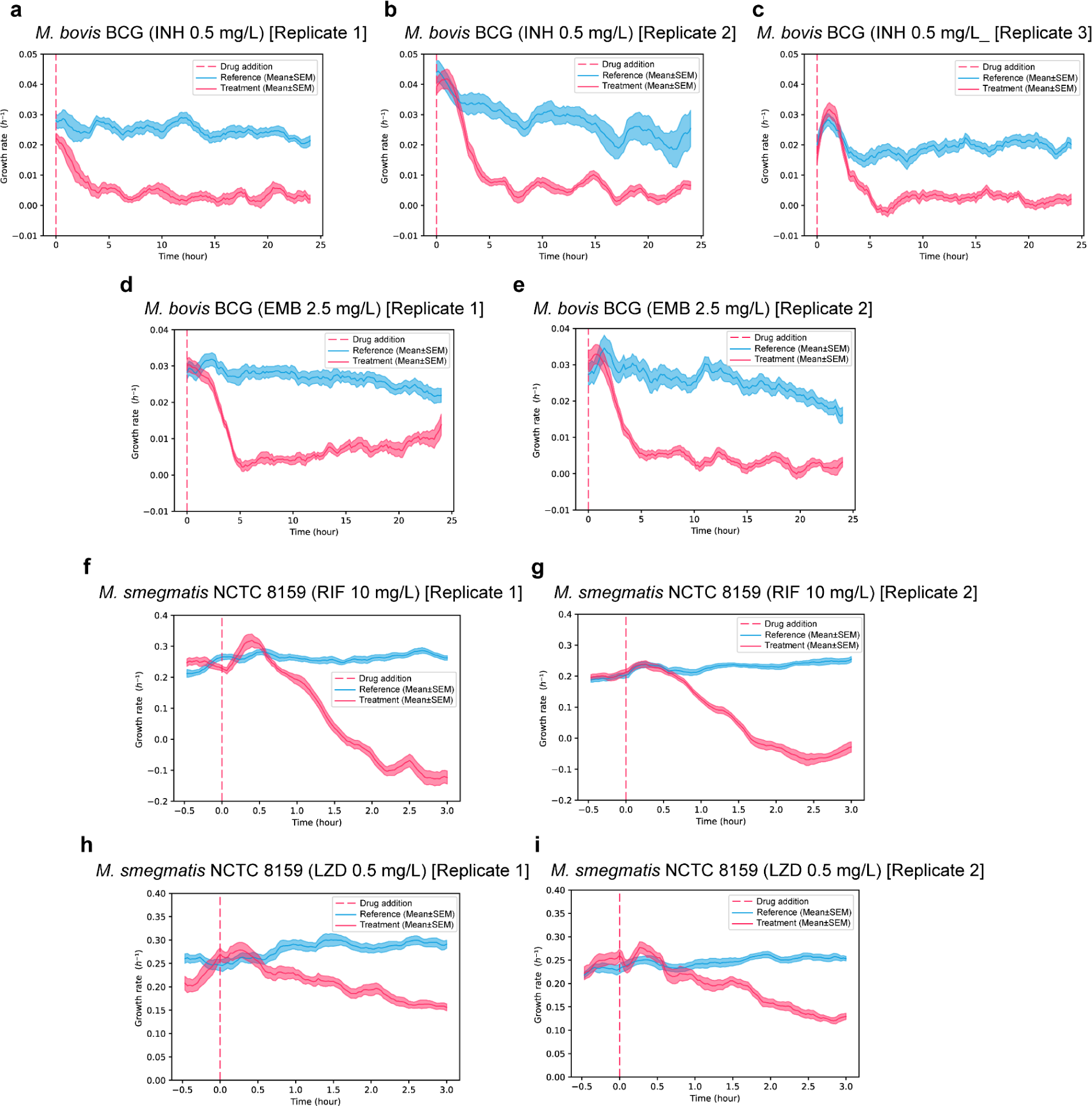
The fast pDST for susceptible mycobacteria in biological replicates. **A - c**. *M. bovis* BCG in isoniazid (INH) 0.5 mg/L. **d - e**. *M. bovis* BCG in ethambutol (EMB) 2.5 mg/L. **f - g**. *M. smegmatis* NCTC 8159 in rifampicin (RIF) 10 mg/L. **h - i**. *M. smegmatis* NCTC 8159 in linezolid (LZD) 0.5 mg/L.

**Supplementary Figure 2.**
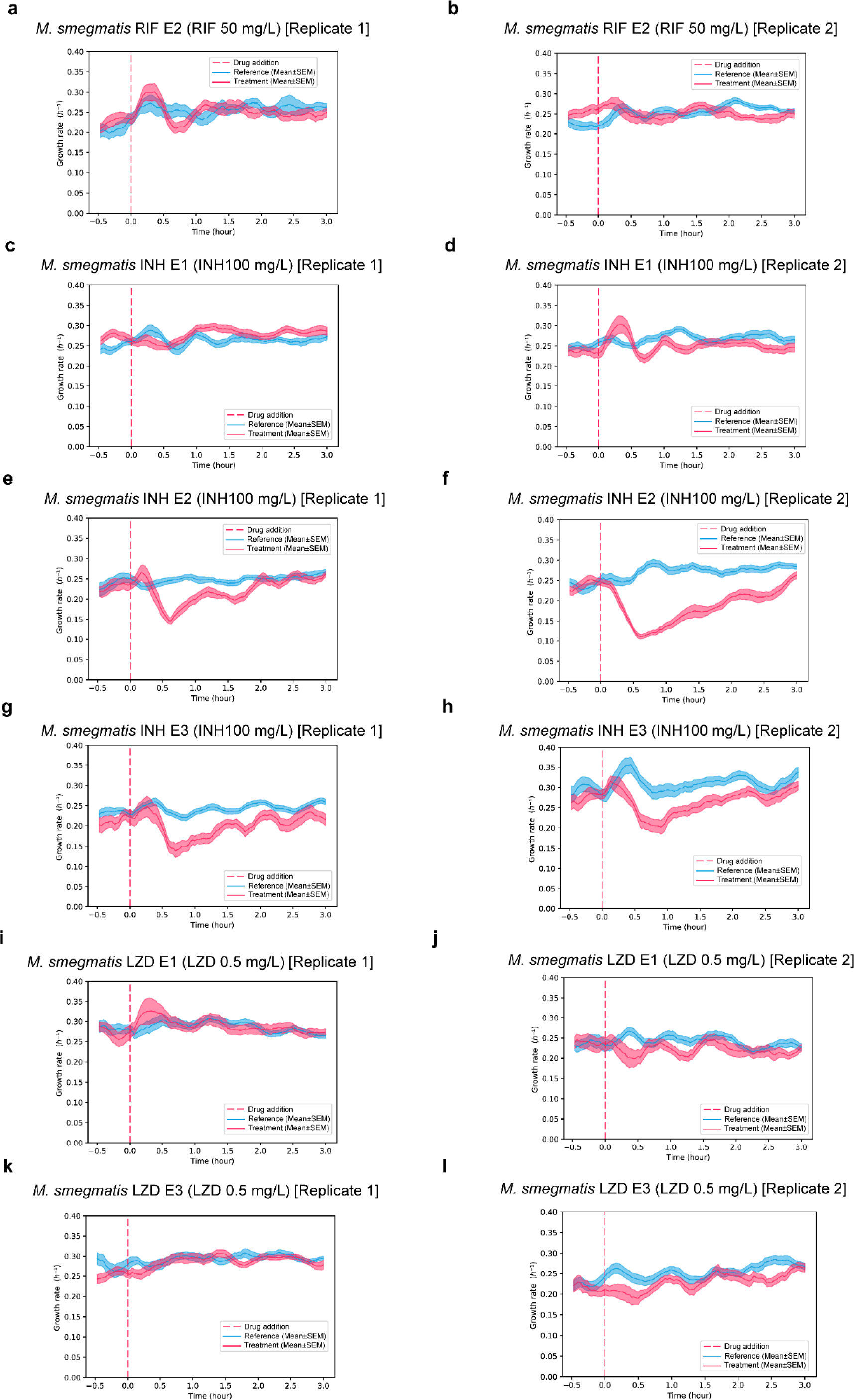
The fast pDST for resistant strains of *M. smegmatis* NCTC 8159 in two biological replicates. **a - b**. *M. smegmatis* NCTC 8159 RIF E3 in rifampicin (RIF) 50 mg/L. **c - d**. *M. smegmatis* NCTC 8159 INH E1 in isoniazid (INH) 100 mg/L. **e - f**. *M. smegmatis* NCTC 8159 INH E2 in isoniazid (INH) 100 mg/L. **g - h**. *M. smegmatis* NCTC 8159 INH E3 in isoniazid (INH) 100 mg/L. **i - j**. *M. smegmatis* NCTC 8159 LZD E1 in linezolid (LZD) 0.5 mg/L. **k - l**. *M. smegmatis* NCTC 8159 LZD E3 in linezolid (LZD) 0.5 mg/L.

